# A novel method for quantitation of AAV genome integrity and residual DNAs using duplex digital PCR

**DOI:** 10.1101/2023.07.19.546797

**Authors:** Lauren Tereshko, Xiaohui Zhao, Jake Gagnon, Tinchi Lin, Trevor Ewald, Marina Feschenko, Cullen Mason

## Abstract

Recombinant adeno-associated virus (rAAV) vectors have become a reliable strategy for delivering gene therapies. As rAAV capsid content is known to be heterogeneous, assays for rAAV characterization are critical for assessing the efficacy and safety of drug products. Multiplex droplet digital PCR (ddPCR) has emerged as a popular molecular approach for characterizing capsid content due to its high level of throughput, accuracy, and replicability. Despite growing popularity, tools to accurately analyze multiplexed data are scarce. Here, we introduce a novel model to estimate genome integrity from duplex ddPCR assays. This work demonstrates that use of a Poisson-multinomial mixture distribution significantly improves the accuracy and quantifiable range of duplex ddPCR assays over currently available models.

## Introduction

rAAV production processes result in heterogeneous populations where in addition to the expected encapsidated full genomes, capsids may be empty, partial, or over-packaged. Unwanted residual DNA from plasmids or host cells involved in the production process may also be encapsidated. Current purification processes can efficiently separate empty from full capsids but cannot differentiate between aberrant versus intended genomes. It is therefore critical to develop assays to thoroughly characterize rAAV encapsidated content and genome integrity. Next generation sequencing (NGS) is an effective tool for characterizing capsid content; however, its utilization is limited for supporting process development due to high material requirements, high cost, and the need for complex data analysis [1–4]. Single and multiplexed digital PCR (dPCR) assays have emerged as effective tools for characterizing encapsidated DNA given their high level of accuracy, low sample requirements and high-throughput capabilities [5,6]. dPCR assays are therefore advantageous for testing drug product and intermediate samples, and for supporting process development of early-stage gene therapy products.

dPCR technologies employ microfluidics to partition template DNA into thousands of independent amplification reactions that are expected to contain zero or one template molecule [7]. End-point fluorescence reactions result in a binary output of partitions that are either negative or positive for template. In singleplex reactions, Poisson statistics can be used to correct for the possibility of multiple templates being partitioned together and can accurately estimate the absolute concentration of template DNA [8,9]. Reactions can be easily duplexed to assess the integrity of individual genes, or rAAV genomes by disrupting capsids prior to template partitioning, and concurrently amplifying targets at the 5’- and 3’- ends of the gene(s) with fluorescent probes of different wavelengths [10,11]. With this strategy, DNA templates containing both targets (double-positive partitions) are considered intact, while templates positive for only one of the targets are considered partial. Unlike singleplex reactions, Poisson statistics are not suitable to model the three-category positive data resulting from duplex reactions [8].

Currently there is a divide between the current technological capability to multiplex assays and the ability to accurately analyze data from such experiments. To bridge the gap, there is a pressing unmet need to develop statistical models that can accurately quantitate end-point fluorescence data from duplex and higher-order multiplexed reactions. Here, we propose a novel Poisson-multinomial model for accurate quantitation of gene and rAAV genome integrity from duplex droplet dPCR (ddPCR) reactions. We compare the accuracy of the model to contemporary statistical models by analyzing duplexed ddPCR data from samples with expected genomes of 0-100% across a range of concentrations. We demonstrate the model expands the dynamic range and improves accuracy of genome integrity estimates compared to simpler models. Finally, we show that integrity values calculated with the Poisson-multinomial model have higher accuracy for both simulated and heat-fragmented rAAV material. These findings establish use of the multinomial Poisson model as a robust approach for analyzing duplex ddPCR data in support of characterizing critical quality attributes of rAAV therapies.

## Results

### I. Comparison of statistical models for genome integrity using plasmid DNA

The accuracy of four analytical models was compared by using mixtures of digested viral vector plasmid (pAAV) to simulate variable degrees of genome integrity. Intact genomes were simulated by restriction enzyme digestion with MfeI outside of the sequence between inverted terminal repeats (ITRs).

Fragmented genomes were simulated by double-digestion outside and between ITR regions (MfeI and NheI) (Fig 1). By combining varying concentration ratios of simulated intact and fragmented genomes, mock samples were produced with either 0, 8, 18, 29, 43, 60, 82 or 100% expected integrities. Duplexed primer/probe sets for 5’ (promoter) and 3’ (poly(A)) genomic regions were used to assess genome integrity of the samples diluted over 12 points to cover the dynamic range of the Bio-Rad QX200 ddPCR system (∼1-5000 copies/µL) [9,12].

**Fig 1.**
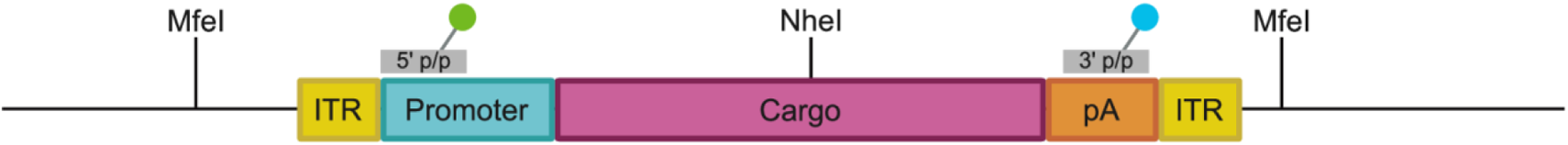
Diagram of pAAV sequence. Restriction sites are indicated by enzyme name and 5’- and 3’- primer/probe sets are indicated by fluorophore illustrations (green and blue respectively).

#### I.I Calculation of genome integrity by simple percentage formula

Bio-Rad’s proprietary ddPCR system outputs the number of droplets in each of four categories (double-positive, single-positive target 1 only, single-positive target 2 only, and empty droplets) and uses a Poisson distribution-based model to calculate the concentration of DNA template molecules in copies per microliter (λ), where p = the fraction of positive droplets in total droplets, and V is the average droplet volume (0.85 nL) (Bio-Rad, personal communication):

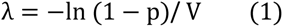

Variations of Formula 1 have been proposed by previous studies to calculate genome integrity as either the percentage of double-positive droplets out of total positive droplets in terms of either droplet number or concentration (Formula 2) [10,13].

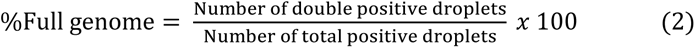

Using a Poisson distribution, it can be determined that when samples are highly dilute (≤150 copies/μL), most positive droplets contain a single DNA template (probability = 0.94), and thereby most double- positive droplets will contain true intact templates as compared to the chance co-localization of 5’ and 3’ targets. As theorized, use of Formula 2 to calculate the genome integrity of the simulated plasmid samples was accurate over only a small portion of the theoretical dynamic range of the QX200 system. Over the tested concentration range (8-5000 copies/µl), for samples with less than 100% expected integrity, the accuracy of calculated genome integrity declined as sample concentration increased, and as expected integrity decreased (Fig 2, Table 1).

**Figure 2.**
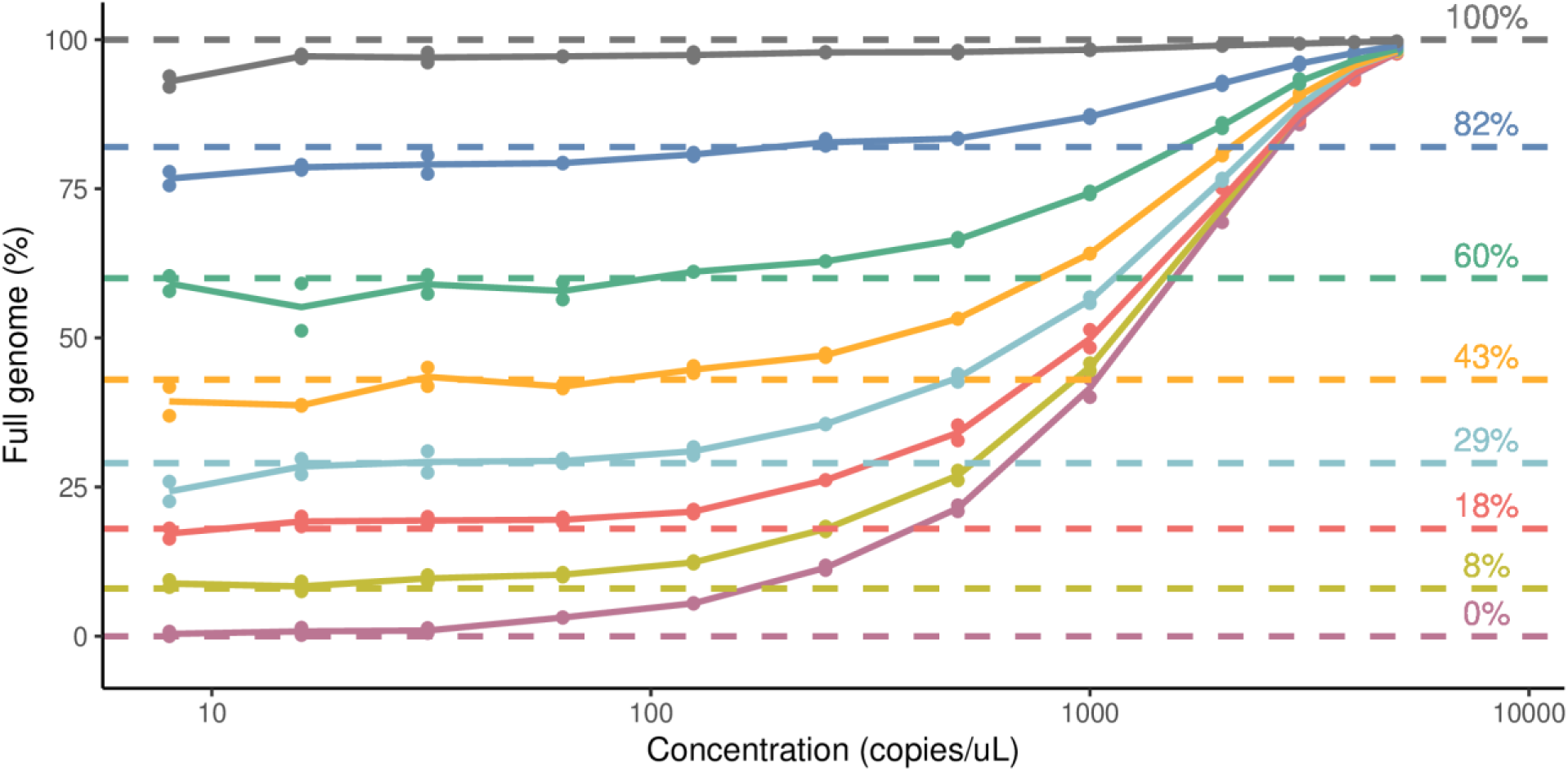
Genome integrity calculated by percentage of double-positive droplets/total positive droplets for mock intact plasmid samples. Genome integrity values calculated by Formula 2 are plotted as single data points (n=1), connecting lines depict average of experimental replicates (N=2). Dashed lines depict the expected integrity values for the samples.

**Table 1.** Percent recovery of genome integrity calculated by percentage of double-positive droplets/total positive droplets for mock intact plasmid samples.

| Copies/μL | Expected genome integrity | | | | | | | |
| --- | --- | --- | --- | --- | --- | --- | --- | --- |
|  | 0% | 8% | 18% | 29% | 43% | 60% | 82% | 100% |
| 8 | NA | 110.2 | 95.5 | 83.6 | 91.5 | 98.5 | 93.6 | 93.0 |
| 16 | NA | 104.3 | 106.7 | 98.0 | 90.0 | 91.9 | 95.8 | 97.2 |
| 31 | NA | 120.8 | 107.6 | 100.7 | 101.1 | 98.3 | 96.4 | 97.0 |
| 63 | NA | 128.6 | 108.3 | 101.3 | 97.3 | 96.4 | 96.7 | 97.2 |
| 125 | NA | 154.3 | 115.9 | 106.9 | 103.9 | 101.8 | 98.5 | 97.4 |
| 250 | NA | 224.5 | 145.2 | 122.5 | 109.5 | 104.7 | 101.0 | 97.9 |
| 500 | NA | 336.6 | 189.3 | 149.3 | 123.8 | 110.8 | 101.7 | 97.9 |
| 1000 | NA | 564.1 | 277.0 | 194.2 | 149.1 | 123.8 | 106.2 | 98.3 |
| 2000 | NA | 898.7 | 407.4 | 263.9 | 188.0 | 142.7 | 113.0 | 99.0 |
| 3000 | NA | 1096.6 | 486.0 | 306.7 | 210.8 | 155.0 | 117.1 | 99.3 |
| 4000 | NA | 1188.6 | 524.7 | 329.2 | 222.4 | 160.8 | 119.2 | 99.6 |
| 5000 | NA | 1224.9 | 544.3 | 337.9 | 228.1 | 164.0 | 120.7 | 99.7 |

Shaded cells indicate recoveries outside of the acceptable range (80-120%).

#### I.II Calculation of genome integrity by physical linkage models

Recently, a formula for genome integrity utilizing the calculation of percent “linkage” has been suggested, where genes contained on the same template are considered linked, and genes that are physically separated are considered unlinked [14]. Linkage is defined as the number of double-positive droplets (linked targets) in excess of what is expected due to chance co-localization of two unlinked targets [15]. The calculation of linked target concentration is included in Bio-Rad’s QuantaSoft raw data file output in terms of copies/µl.

In instances where the concentration of each target is similar, the authors suggest that percent linkage (i.e. genome integrity) can be calculated by dividing the linkage concentration by the average concentrations of the 5’ and 3’ target (abbreviated prom and pA respectively) (Formula 3). The authors note that if the concentrations of the two targets are unequal due to amplification bias resulting from experimental conditions (method-induced genome fragmentation, differences in genome accessibility, or differences in amplicon size) that a compensated version of the equation can be used which involves adjustment of the linkage value by addition of the absolute value of the difference in concentrations of the 5’ and 3’ target, followed by dividing by the maximum value of either the concentration the 5’ or the 3’ target (Formula 4) [14].

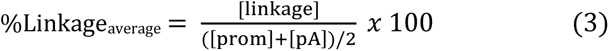

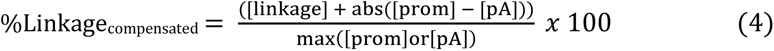

When either Formula 3 or Formula 4 was used to calculate the percent genome integrity of the plasmid mock samples, the results were relatively stable across the tested concentration range (8-5000 copies/µl), as demonstrated by the assessment of the relative standard deviation (RSD) across concentrations. Of note, for both models, the calculated integrity values were significantly more variable as the expected integrity of the samples decreased. Additionally, the integrity values for all fragmented genome samples were overestimated as demonstrated by the calculated recoveries relative to the expected values, and the overestimations became more pronounced as expected integrity decreased (Fig 3 A, B, Table 2). In this data set, the amplification of the 5’ and 3’ targets are even, and the calculated values from Formula 3 and Formula 4 are therefore expected to be similar. Formula 3 was more accurate and more precise than Formula 4 across the full range of integrity (Table 2). The lack of accuracy and precision of Formula 3 for samples with lower integrity would require shortening the linear range of the method (Fig 3 A, B, D).

**Fig 3.**
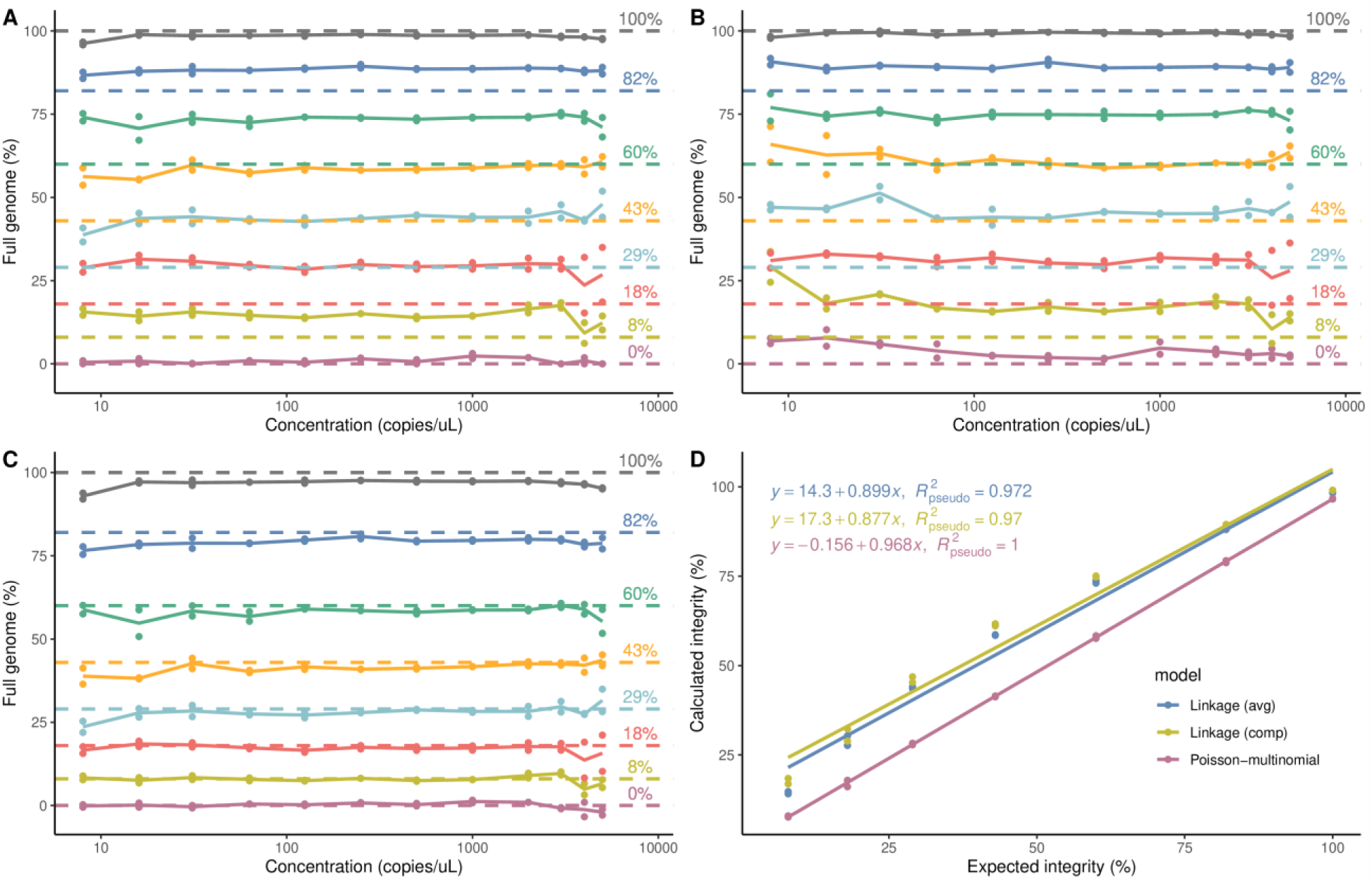
Genome integrity calculated by linkage and Poisson-multinomial models for mock intact plasmid samples. A), average percent linkage (Formula 3), B), compensated percent linkage (Formula 4). C), Poisson-multinomial (Formula 7). Calculated percent integrity values are plotted as single data points (n=1), connecting lines depict average of experimental replicates (N=2). Dashed lines depict the expected values for the samples. D), Linearity of models. Average calculated percent integrity values from sample dilutions are plotted as points (n=12, N=2).

**Table 2.**
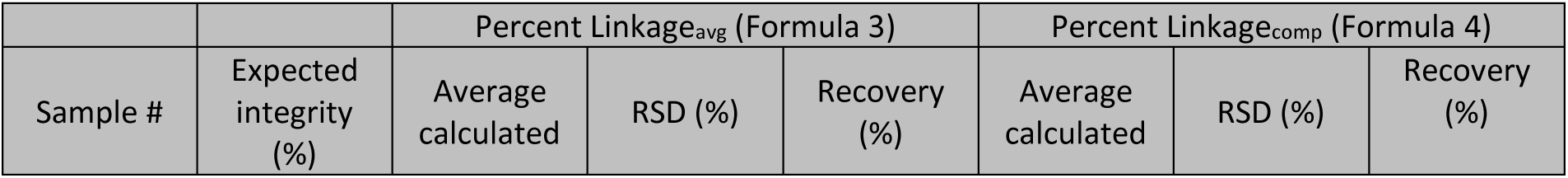
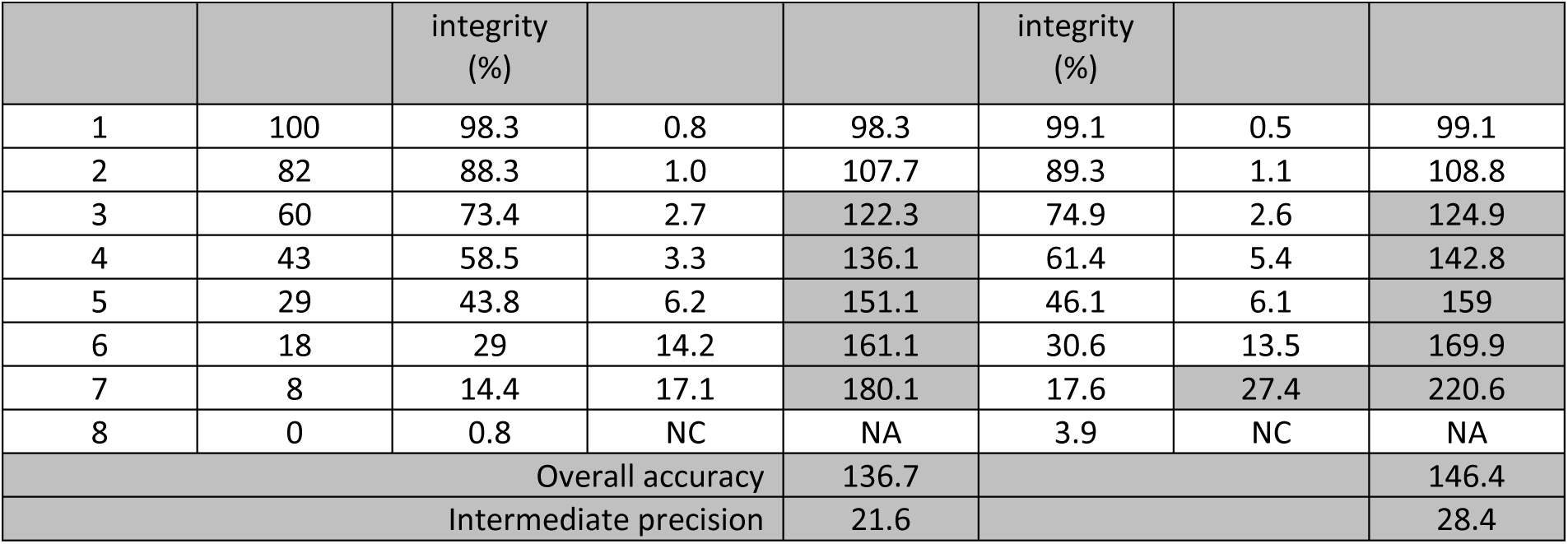
Genome integrity and recovery calculated by linkage models.

Each sample was tested at twelve dilutions and the results were then averaged to produce a calculated integrity value and percent RSD. Shaded cells indicate recoveries outside of the acceptable range (80-120%) or RSD greater than 20%. RSD not calculated for 0% expected integrity (NC).

#### I.III Calculation of genome integrity by Poisson-multinomial model

As an alternative method for genome integrity calculation, we developed a more robust statistical model that utilizes a Poisson-multinomial mixture distribution. In the model below, three categories of positive droplets are considered with *k* molecules as a multinomial model with three species containing 1.) 5’ target only, 2.) 3’ target only, and 3.) both 5’ and 3’ targets (double-positive).

To model the probability of *k* molecules within a droplet, we can utilize the binomial distribution, *p(k)* with *k* successes and *m* trials, where *m* is the total number of molecules [16]. However, when the number of droplets is large, the binomial distribution can be approximated by the Poisson distribution with parameter, lambda.

This Poisson-multinomial model has two unknown parameters, *pprom* and *ppolyA*. To estimate these parameters, the measured numbers of promoter single-positive droplets, poly(A) single-positive droplets, and the total number of droplets are needed. The upper limit of the dynamic range of the QX200 ddPCR plate reader is 5000 copies/μL. Given that the average droplet volume is 0.85 nL, this upper limit is equal to 4.25 copies/droplet. It can be further determined by the Poisson distribution that the probability of more than 20 copies of template molecules being present in a droplet is negligible (probability = 5.4E−9). It is therefore numerically sufficient to model the situation where the average number of template molecules within a droplet is ≤ 20 copies.

The calculated number of promoter single positive-droplets across all droplets can be derived from the Poisson-multinomial model where *k* = the number of template molecules within a droplet, and *D* is the total number of droplets (Formula 5).

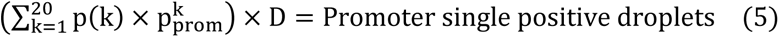

Similarly, the calculated number of poly(A) single-positive droplets across all droplets can be derived as Formula 6.

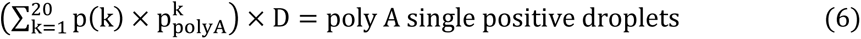

Solving the 20^th^ degree polynomial gives us the unknown quantities *pprom* and *ppolyA*. Genome integrity can then be calculated as the expected (E) number of full genomes across all droplets divided by the expected number of total DNA template molecules across total droplets:

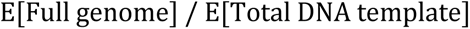

where the numerator quantity is derived from the double expectation rule:

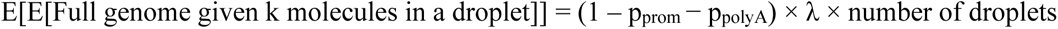

and the denominator is given by:

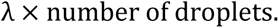

With these expressions, Formula 7 can be used to calculate the percentage of templates that are fully intact:

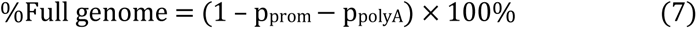

When the Poisson-multinomial model was used to calculate the percent genome integrity of the plasmid mock samples, all values were consistent (RSD <20%) and aligned with the expected values (recoveries between 80-120%) across the tested concentration range (8-5000 copies/µL) (Fig 3C, Table 3). The overall accuracy of the Poisson-multinomial distribution model was 96.3% and the accuracy was consistent across the full range of intact genomes (0-100%). Although the calculated integrity values were significantly more variable as the integrity of the mock plasmid samples decreased, the Poisson- multinomial model was more precise than linkage models (intermediate precision <1%), as calculated by the RSD of all the recoveries (Table 3). Plotting the sample integrity versus the calculated integrity values for each model shows that the Poisson-multinomial model is more linear than linkage models throughout the range of possible values (0-100%) when the data were fit using generalized least squares (GLS) to account for correlations among experimental replicates (Fig 3D, S1 Fig, S1 Table).

**Table 3.** Genome integrity calculated by the Poisson-multinomial distribution model.

| Sample # | Expected integrity (%) | Average calculated integrity (%) | RSD (%) | Recovery (%) |
| --- | --- | --- | --- | --- |
| 1 | 100 | 96.7 | 1.4 | 96.7 |
| 2 | 82 | 79.1 | 1.7 | 96.4 |
| 3 | 60 | 58 | 4.1 | 96.6 |
| 4 | 43 | 41.4 | 4.8 | 96.2 |
| 5 | 29 | 28 | 8.2 | 96.7 |
| 6 | 18 | 17 | 15.8 | 94.3 |
| 7 | 8 | 7.8 | 18.0 | 96.9 |
| 8 | 0 | -0.1 | NC | NA |
| Overall accuracy |  |  |  | 96.3 |
| Intermediate precision |  |  |  | 0.9% |

Each sample was tested at twelve dilutions and the results were then averaged to produce a calculated integrity value and percent RSD. Shaded cells indicate recoveries outside of the acceptable range (80- 120%) or RSD greater than 20%. RSD not calculated for 0% expected integrity (NC).

### II. Comparison of Linkage vs Poisson-multinomial models for mock genome integrity using AAV material

To evaluate the Poisson-multinomial model for genome integrity empirically with rAAV material, fragmented rAAV genomes were simulated by combining two samples containing common promoter sequences but differing poly(A) sequences, in varying concentration ratios. Primer/probe sets were designed to target the shared promoter region of Samples 1 and 2, and the poly(A) sequence of Sample 1 (Fig 4). Sample 1 was first assayed with both primer/probe sets in a duplex reaction to estimate genome integrity from dilutions <250 copies/µl using Formula 2. The calculated integrity of Sample 1 (84%) was then set as the maximal expected integrity value. When assayed with the same primer/probe sets, Sample 2 is expected to have no amplification of the poly(A) sequence (0% intact). In this manner, six mock integrity samples were prepared with expected integrities of either 0, 5,10,21,42,63 or 84%.

**Fig 4.**
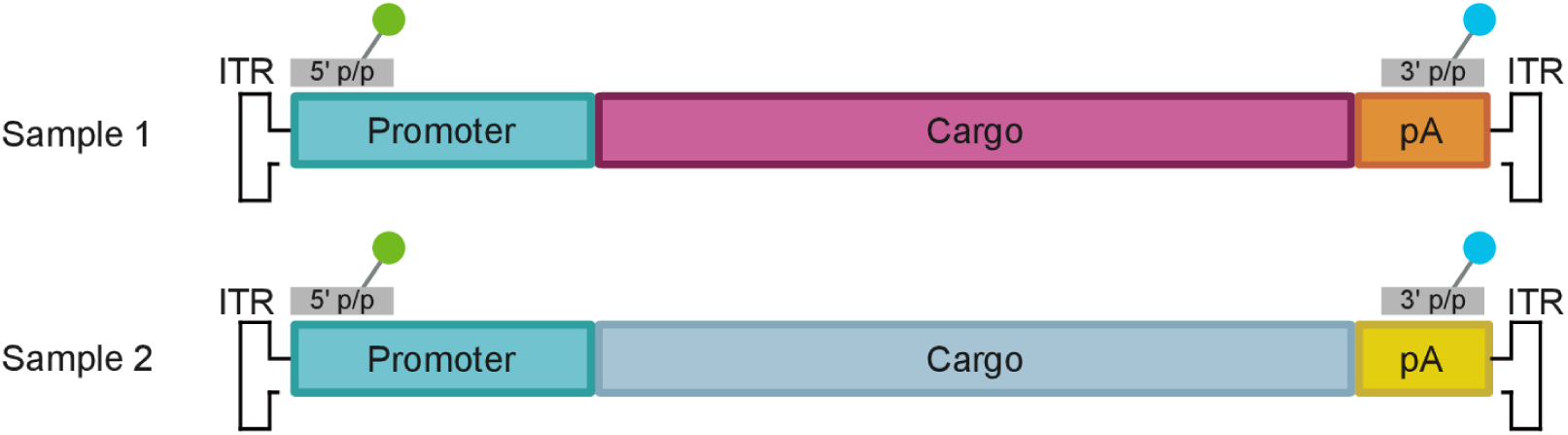
Diagram of AAV sequences. Sample 1 and sample 2 genomes share a common promoter but differ in poly(A) sequence. 5’- (green) and 3’- (blue) primer/probe sets are indicated by fluorophore illustrations.

It is possible that rAAV samples may contain both targets within a single capsid without the targets being physically linked. To truly assess genome integrity as opposed to double-positive capsids, the samples must be decapsidated prior to droplet generation. Capsid serotypes differ in their thermal stabilities, and thermal lysis protocols with high heat have been shown to reduce genome integrity due to increased rates of spontaneous hydrolysis of the phosphate DNA backbone [14,17–19]. Decapsidation was therefore performed by alkaline lysis and a short incubation at low heat (10 minutes, 60°C) to minimize hydrolysis. Duplexed assays were performed as described previously for plasmid samples; however, samples were tested over four dilutions that spanned a narrower range of concentrations (62- 498 copies/µl).

The genome integrity of each sample was calculated using either average linkage percentage (Formula 3) or the Poisson-multinomial model. Comparison with the compensated linkage percentage (Formula 4) was not included as the concentrations of promoter and poly(A) are expected to be uneven due to the experimental design. Formula 3 yielded consistent results across the range of concentrations tested; however, the integrity values were consistently overestimated for all samples (Fig 5A, Table 4). The Poisson-multinomial model yielded consistent results across the range of concentrations tested with higher accuracy (Fig 5B, Table 4).

**Fig 5:**
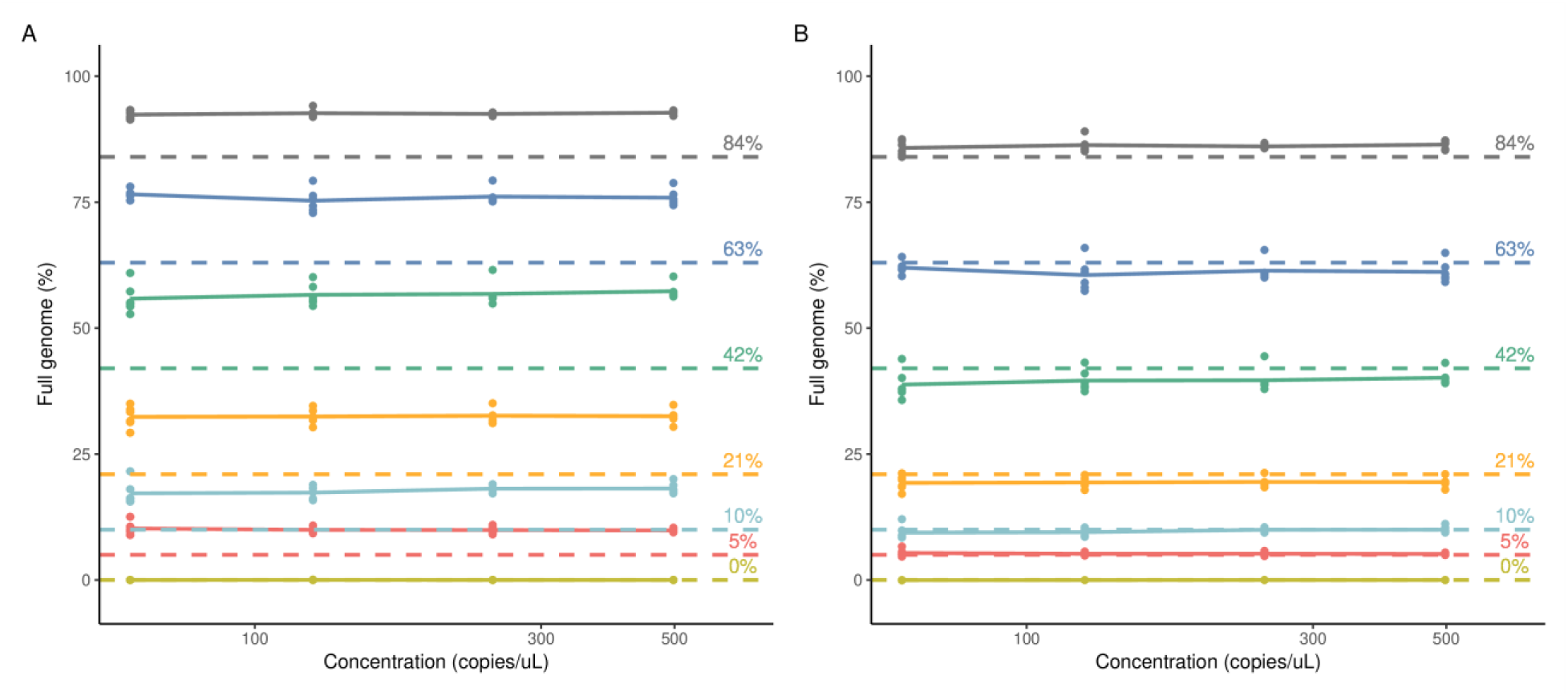
Comparison of mock rAAV genome integrities. Calculated with A.) average percent linkage model (Formula 3) or B.) Poisson-multinomial model. Calculated percent integrity values are plotted as single data points (n=1), connecting lines depict average of experimental replicates (N=6). Dashed lines depict the expected values for the samples.

**Table 4.** Comparison of average AAV genome integrities and percent recovery calculated with average percent linkage model vs Poisson-multinomial.

| Sample # | Expected integrity | Percent Linkage<sub>avg</sub> |  |  | Poisson-multinomial |  |  |
| --- | --- | --- | --- | --- | --- | --- | --- |
|  |  | Avg. replicate integrity (%) | RSD (%) | Recovery (%) | Avg. replicate integrity (%) | RSD (%) | Recovery (%) |
| 1 | 84 | 93.4 | 0.6 | 111.1 | 87.6 | 1.1 | 104.3 |
|  |  | 92.3 | 0.8 | 109.9 | 85.5 | 1.3 | 101.8 |
|  |  | 92.2 | 0.6 | 109.7 | 85.6 | 1.1 | 101.9 |
|  |  | 92.5 | 0.5 | 110.1 | 86.2 | 0.9 | 102.7 |
|  |  | 92.7 | 0.4 | 110.3 | 86.3 | 0.8 | 102.7 |
|  |  | 92.4 | 0.4 | 110.0 | 85.8 | 0.8 | 102.1 |
| 2 | 63 | 78.9 | 0.7 | 125.2 | 65.1 | 1.2 | 103.4 |
|  |  | 76.3 | 0.3 | 121.1 | 61.6 | 0.7 | 97.8 |
|  |  | 75.0 | 1.4 | 119.0 | 60.0 | 2.2 | 95.3 |
|  |  | 75.0 | 2.2 | 119.0 | 60.0 | 3.4 | 95.2 |
|  |  | 75.2 | 1.1 | 119.4 | 60.2 | 1.8 | 95.6 |
|  |  | 75.5 | 1.1 | 119.9 | 60.7 | 1.9 | 96.4 |
| 3 | 42 | 60.7 | 1.1 | 144.6 | 43.6 | 1.4 | 103.9 |
|  |  | 56.0 | 4.2 | 133.4 | 38.9 | 5.8 | 92.5 |
|  |  | 55.6 | 1.7 | 132.4 | 38.5 | 2.2 | 91.6 |
|  |  | 55.5 | 2.0 | 132.2 | 38.4 | 2.5 | 91.4 |
|  |  | 56.2 | 1.5 | 133.8 | 39.0 | 2.1 | 92.9 |
|  |  | 55.8 | 1.9 | 132.9 | 38.8 | 2.8 | 92.3 |
| 4 | 21 | 34.8 | 0.7 | 165.9 | 21.1 | 0.8 | 100.5 |
|  |  | 31.6 | 5.1 | 150.4 | 18.7 | 6.0 | 89.2 |
|  |  | 30.8 | 1.7 | 146.7 | 18.2 | 1.8 | 86.6 |
|  |  | 32.0 | 1.4 | 152.6 | 19.1 | 2.0 | 90.9 |
|  |  | 32.7 | 1.5 | 155.6 | 19.5 | 1.6 | 92.9 |
|  |  | 33.0 | 2.6 | 157.1 | 19.7 | 2.8 | 94.0 |
| 5 | 10 | 19.7 | 7.3 | 197.3 | 10.9 | 8.1 | 109.4 |
|  |  | 18.1 | 6.8 | 180.7 | 9.9 | 7.4 | 99.4 |
|  |  | 17.2 | 6.4 | 172.0 | 9.4 | 7.1 | 94.1 |
|  |  | 17.0 | 5.3 | 170.1 | 9.3 | 5.8 | 92.8 |
|  |  | 17.5 | 5.4 | 174.6 | 9.6 | 5.6 | 95.7 |
|  |  | 16.9 | 6.4 | 168.6 | 9.2 | 7.2 | 91.9 |
| 6 | 5 | 11.2 | 8.5 | 223.6 | 5.9 | 9.0 | 117.8 |
|  |  | 10.0 | 3.8 | 200.5 | 5.3 | 4.0 | 105.5 |
|  |  | 9.7 | 3.9 | 194.9 | 5.1 | 3.5 | 102.0 |
|  |  | 9.6 | 6.7 | 192.7 | 5.1 | 7.4 | 101.1 |
|  |  | 9.8 | 6.5 | 196.8 | 5.2 | 6.7 | 103.4 |
|  |  | 9.5 | 1.4 | 190.3 | 5.0 | 1.3 | 99.5 |
| 7 | 0 | 0.0 | NC | NA | NA | NC | NA |
|  |  | 0.0 | NC | NA | NA | NC | NA |
|  |  | 0.0 | NC | NA | NA | NC | NA |
|  |  | 0.0 | NC | NA | NA | NC | NA |
|  |  | 0.0 | NC | NA | NA | NC | NA |
|  |  | 0.0 | NC | NA | NA | NC | NA |
| Overall accuracy |  |  |  | 149.6 |  |  | 98.1 |
| Intermediate precision |  |  |  | 21.8 |  |  | 6.5 |

Highlighted cells >120% recovery. RSD not calculated for 0% expected integrity (NC).

To compare the accuracy and precision of the models, the percent recoveries of calculated genome integrities were evaluated in six independent experiments. When integrity was calculated using the average percent linkage model, all sample replicates showed acceptable precision (RSD<20%), however samples <84% expected integrity over-recovered (>120%). As seen with plasmid mock samples, over- estimation of integrity worsened as expected integrity decreased. In comparison, when the Poisson-multinomial model was used, all sample replicates demonstrated acceptable recovery (between 80- 120%) and precision (RSD<20%), as summarized in Table 4. Compared to the average percent linkage model, the Poisson-multinomial model showed greater overall accuracy and intermediate precision (149.6 vs 98.1%, and 21.8 vs 6.5% respectively).

To compare the linearity of the models, the average calculated percent genome integrities of each sample replicate (summarized in Table 4) were plotted against the expected integrity values, and the data were fit using GLS to account for correlations among experimental replicates. Although both models showed linear sample recovery in the range of 5-84%, the Poisson-multinomial data had a better linear fit with a slope of 1.01 and pseudo R^2^ of 0.996 (Fig 6, S2 Fig, S2 Table).

**Fig 6.**
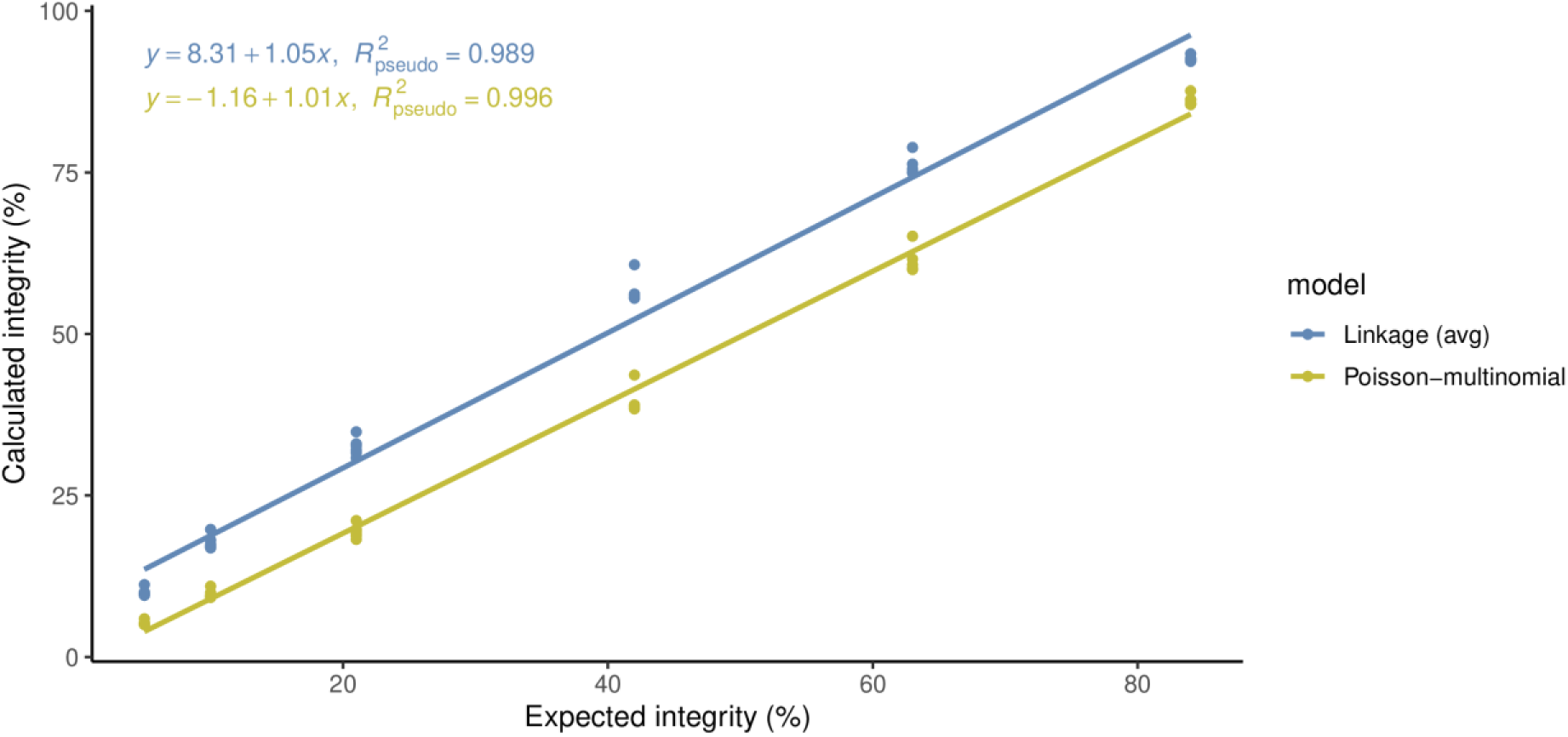
Linearity of the Poisson-multinomial model for mock rAAV genome integrity. Average calculated percent integrity values from experimental replicates are plotted as points (n=4, N=6).

### III. Comparison of Linkage vs Poisson-multinomial models for genome integrity of fragmented rAAV material

To evaluate the Poisson-multinomial model in a more realistic context, heat-fragmented genomes were prepared as thermal stress is known to degrade DNA [20,21]. Replicate rAAV samples were decapsidated using the standard protocol followed by either a 0, 1, 5, 10, 20, or 30-minute incubation at 95°C. Primer/probe sets targeting the promoter and poly(A) sequences were used to assay the samples in duplex reactions. Sample material was previously determined to have an integrity of 48% via long-read NGS.

As seen in Fig 7, although genome integrity decreased as incubation time at 95°C increased as expected for all models, the values were overestimated when calculated by either linkage-based model compared to the Poisson-multinomial model.

**Fig 7.**
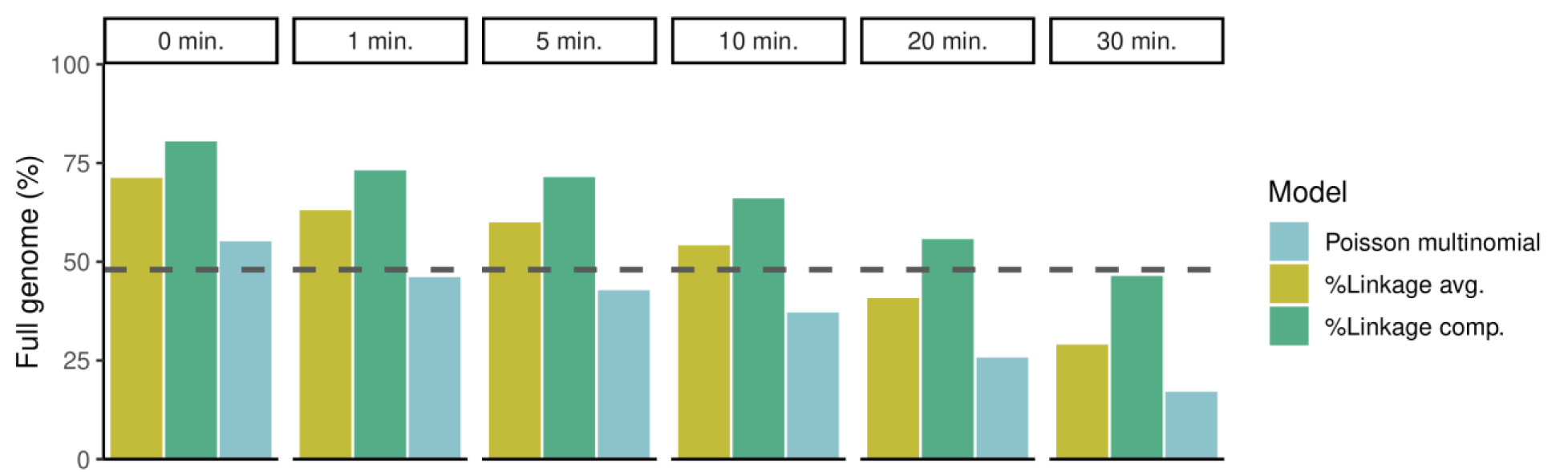
Genome integrities of heat-fragmented rAAV calculated by linkage or Poisson-multinomial models. Average calculated genome integrity values are plotted as single bars (n=12, N=2). Dashed line depicts expected integrity as determined by long-read NGS.

## Discussion

This work summarizes the development of a novel analytical model for the calculation of gene and genome integrity from duplexed ddPCR assays. The new model has been coded in R as a Shiny application, and the code is publicly available. Use of the Poisson-multinomial mixture distribution offers more rigorous modeling compared to simplistic percentage-based calculations (described by Formula 2), which are accurate only at highly dilute concentrations. Comparatively, our model is highly accurate across a wide range of template concentrations, expanding the dynamic concentration range of duplex integrity assays to at least four orders.

Unlike Poisson models which are limited to the prediction of single categories of droplet species (positive or negative), the Poisson-multinomial model considers the probability of multiple species of positive droplets, making it more suitable for the interpretation of duplexed data. Although linkage models also consider multiple species of positive droplets by subtracting the number of droplets with linked targets expected by chance from the number of double-positive droplets, we found our model to have improved intermediate precision and accuracy (compared to Formulas 3 and 4). Critically, use of linkage models consistently resulted in inflated integrity values for both simulated integrity samples (plasmid and rAAV) and heat-degraded rAAV samples.

We speculate linkage models may over-estimate genome integrity by calculating the fraction of full genomes (concentration of linked targets) out of the concentration of genome fragments (as either the average or maximum of 5’ and 3’ target concentrations) when the total concentration of both targets (sum of 5’ and 3’ target concentrations, minus linked target concentration) should be considered instead. The linkage models may have been developed with the assumption that even if genomes are fragmented that both 5’ and 3’ regions will be present, when in fact, truncated genomes often have fragmentation bias. Capsids containing truncated rAAV genomes can result from a variety of reasons such as defective replication or errors in viral packaging [22–24]. Encapsidation errors have been shown to have bias towards 5’ truncation [24,25]. Furthermore, the type and rate of truncation event is also dependent on the production method, vector sequence, and whether the virus is single-stranded or self-complementary [26–28]. For these reasons, we believe it is more appropriate to consider the total concentration of template DNA as opposed to the average (or maximum) of 5’ and 3’ targets.

Accurately characterizing the encapsidated DNA of gene therapy material is a critical aspect of drug product safety and efficacy. Although several orthogonal approaches should be combined to completely characterize capsid content (dPCR, NGS, TEM, AUC, functional potency, etc.), duplexed ddPCR assays offer the potential to increase throughput and decrease the cost of molecular characterization. By expanding the concentration dynamic range, as well as expanding the linear range of the integrity values to cover >0 to 100%, the Poisson-multinomial model is well suited to serve as a screening method to support DOE studies and guide process decisions.

The work summarized here for quantitating genome integrity can be applied to other duplexed ddPCR assays, including those aimed at monitoring residual DNA size and identity. FDA guidance recommends that residual DNAs be limited to under 10 ng/dose and less than 200 base pairs in length in final drug product (Yang 2013). As contaminants are expected to be present in very low concentrations in final drug products, utilizing the Poisson-multinomial model for accurate analysis of residual DNA integrity may therefore be a useful tool in assessing the integrity and therefore risk level of contaminant DNAs. Multiplexed ddPCR methods could potentially be expanded beyond duplex reactions to simultaneously measure both the quantity and integrity of specific residual DNAs from plasmids or host cell genes. By controlling whether decapsidation occurs prior to, or after droplet generation, one could potentially determine the levels of encapsidated residuals relative to the rAAV genome and provide characterization data around different capsid populations: those that contain the rAAV genome and those that do not. Such analyses would require both the viral genome and residual plasmids to be detectable at similar dilutions, for which the expanded dynamic range of the Poisson-multinomial model may be advantageous.

## Materials and Methods

### ddPCR

All samples were partitioned into approximately 20,000 droplets using a Bio-Rad Automated Droplet Generator (1864101). Droplets were subjected to ITR restriction digestion and endpoint PCR thermal cycling and read on a Bio-Rad QX200 droplet reader with QuantaSoft software (version 1.7.4).

### Sample preparation

#### Plasmid simulated genomes

pAAV was digested with MfeI alone or with MfeI and NheI together. DNA concentrations of digests were measured using a Nanodrop spectrophotometer and converted to copies/ml. Digested plasmids (MfeI and MfeI/NheI) were diluted in TE and mixed at varying ratios to simulate varying degrees of genome fragmentation.

#### rAAV simulated genomes

Mock integrity samples were prepared by mixing varying ratios of two rAAVs containing the same promoter and different poly(A) tails. Samples were pre-diluted to a target concentration range (1.125e5- 5e6 vg/µL) and digested with DNase I for 30 minutes at 37°C. Viral capsids were disrupted with SDS solution and incubation at low heat (10 minutes, 60°C). Decapsidated samples were serially diluted and combined with ddPCR master mixes in 96-well PCR plates.

#### Heat fragmented rAAV genomes

rAAV samples were pre-diluted, DNase treated and decapsidated following the procedure described for rAAV simulated genomes. Following decapsidation, samples were incubated at 95°C for either 0, 1, 5, 10, 20 or 30 minutes and then serially diluted and combined with ddPCR master mixes in 96-well PCR plates.

#### Long-read NGS

Independent preparations of rAAV were sequenced on two instruments: MinION Mk1B (Oxford Nanopore Technologies) and Sequel II (PacBio). Percent genome integrity was estimated using a custom analytical pipeline. Genome integrity results from both sequencers were averaged and used as the expected integrity for AAV genome in heat fragmentation experiments.

### Data analysis

Droplet analysis was performed using BioRad QuantaSoft software (version 1.7.4). Inter-assay replicates and experimental replicates are indicated in the figure legends. Percent recovery was calculated by comparing the measured genome integrity to the expected genome integrity using the following

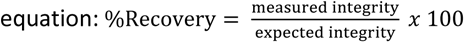

Percent relative standard deviation (RSD) was calculated as the standard deviation of the calculated integrity replicates divided by the average of the calculated integrity replicates, multiplied by 100. Assay accuracy was calculated as the grand average of the replicate average percent recovery values.

Intermediate precision was calculated as the RSD of the replicate average percent recovery values.

### Shiny app for Poisson multinomial model

A Web-based Shiny application was developed and deployed on the cloud-based R/Posit platform[22,23].

#### Shiny app description

Data files generated by the QX200 droplet reader are imported into the app which extracts relevant raw data to calculate genome integrity with the Poisson multinomial model. Each well in the assay plate has two rows which correspond to two respective targets. Columns B through L are raw data extracted from the raw data file, while genome integrity is appended in column M in duplicate rows for each well. It is worth noting that “NA” will be displayed in column M when any droplet number in columns I-K is 0, as it fails to meet the prerequisites for Poisson-multinomial model. To ensure data integrity, the application verifies that the two percent full genome values in duplicate rows are identical, which is displayed as Boolean values in column N.

## Supporting information

Supporting_info_Tereshko_etal_2023

## Acknowledgements

We thank Biogen’s NGS & Genetic Technologies Lab for assistance with sequencing rAAV material. We thank Romi Admanit and Svetlana Bergelson for analytical review and resource management.

