## Supporting_info_Tereshko_etal_2023 for "A novel method for quantitation of AAV genome integrity and residual DNAs using duplex digital PCR"

**Supporting information**

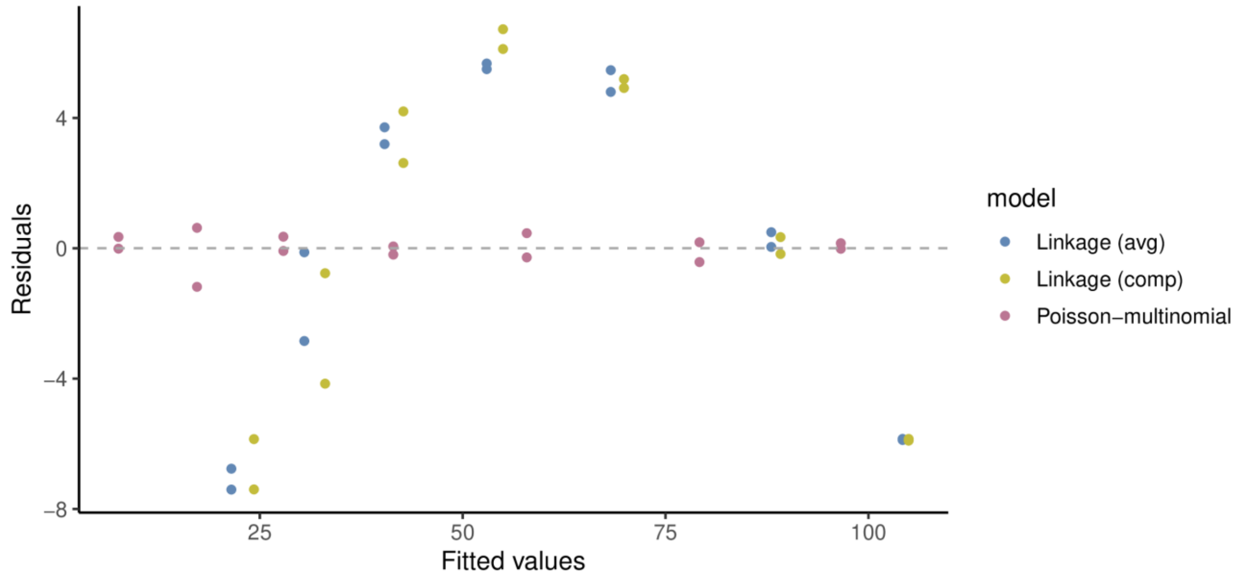


**S1 Fig. Linearity of Linkage and Poisson-multinomial models for plasmid genome integrity.** Fitted values vs residuals plotted for each model.

**S1 Table. Linear fit information for linear models plotted in Figure S1.**

| | | Coefficient | Standard error | p-value |
|---|---|---|---|---|
| Linkage(avg) (pseudo R2 =0.972) | Intercept | 14.3 | 3.89 | 0.003 |
| | Slope | 0.899 | 0.67 | <0.001 |
| Linkage(comp) (pseudo R2 =0.970) | Intercept | 17.3 | 3.98 | 0.001 |
| | Slope | 0.877 | 0.07 | <0.001 |
| Poisson-multinomial (pseudo R2 1.000) | Intercept | -0.156 | 0.12 | 0.23 |
| | Slope | 0.968 | 0.002 | <0.001 |

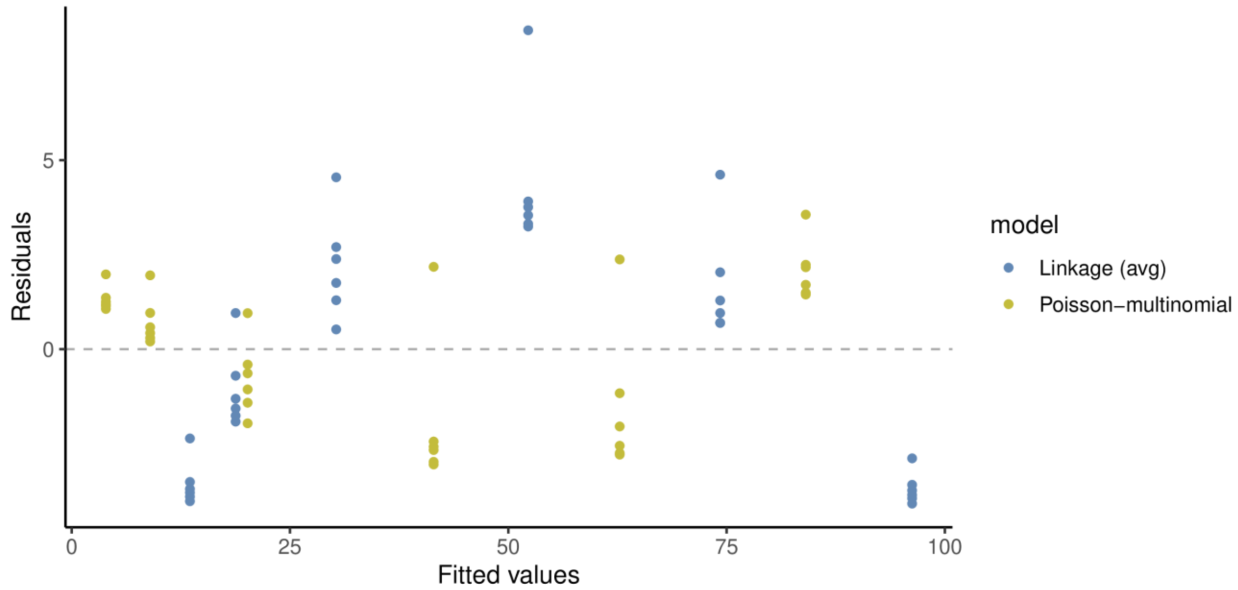


**S2 Fig. Linearity of Linkage and Poisson-multinomial models for rAAV genome integrity.** Fitted values vs residuals plotted for each model.

**S2 Table. Linear fit information for linear models plotted in Figure S2.**

| | | Coefficient | Standard error | p-value |
|---|---|---|---|---|
| Linkage(avg) (pseudo R2 =0.989) | Intercept | 8.31 | 2.48 | 0.002 |
| | Slope | 1.05 | 0.05 | <0.001 |
| Poisson-multinomial (pseudo R2 0.996) | Intercept | -1.16 | 0.59 | 0.058 |
| | Slope | 1.01 | 0.01 | <0.001 |
